# Different genetic architectures of complex traits and their relevance to polygenic score performance in diverse populations

**DOI:** 10.1101/2022.10.29.514295

**Authors:** Nuno R. G. Carvalho, Adrian M. Harris, Joseph Lachance

**Affiliations:** School of Biological Sciences, Georgia Institute of Technology, Atlanta, GA, USA

**Keywords:** complex traits, genetic architecture, Gini coefficients, polygenic scores, population genetics, portability

## Abstract

**Background:** Despite the many insights gleaned from GWAS, polygenic predictions of complex traits have had limited success, particularly when these predictions are applied to individuals of non-European descent. A deeper understanding of the genetic architecture of complex traits may inform why some traits are easier to predict than others.

**Methods:** Examining 163 complex traits from the UK Biobank, we compared and contrasted three aspects of genetic architecture (SNP heritability, LD variability, and genomic inequality) with three aspects of polygenic score performance (prediction accuracy in the source population, portability across populations, and trait divergence across populations). Here, genomic inequality refers to how unequally the genetic variance of each trait is distributed across the top trait-associated SNPs, as quantified via a novel application of Gini coefficients.

**Results:** Consistent with reduced statistical power, polygenic predictions of binary traits performed worse than predictions of quantitative traits. Traits with low Gini coefficients (i.e., highly polygenic architectures) include hip circumference as well as systolic and diastolic blood pressure. Traits with large population-level differences in polygenic scores include skin pigmentation and hair color. Focusing on 96 quantitative traits, we found that highly heritable traits were easier to predict and had predictions that were more portable to other ancestries. Traits with highly divergent polygenic score distributions across populations were less likely to have portable predictions. Intriguingly, LD variability was largely uninformative regarding the portability of polygenic predictions. This suggests that factors other than the differential tagging of causal SNPs drive the reduction in polygenic score accuracy across populations. Subsequent analyses identified suites of traits with similar genetic architecture and polygenic score performance profiles. Importantly, lifestyle and psychological traits tended to have low heritability, as well as poor predictability and portability.

**Conclusions:** Novel metrics capture different aspects of trait-specific genetic architectures and polygenic score performance. Our findings also caution against the application of polygenic scores to traits like general happiness, alcohol frequency, and average income, especially when polygenic scores are applied to individuals who have an ancestry that differs from the original source population.

## Background

Recent years have seen an explosive growth in our understanding of the genetics of complex traits [1–3]. Large datasets, such as the UK Biobank [4], have facilitated the genetic analysis of complex traits, and to date over 6400 genome-wide association studies (GWAS) have yielded over 529,000 associations [5]. Despite these discoveries, major knowledge gaps still exist [6]. How can the field go beyond mere catalogs of statistical associations? One way that GWAS results can be leveraged is to infer the genetic architecture of complex traits. A second downstream application of GWAS involves generating polygenic predictions of complex traits and hereditary disease risks.

Genetic architecture refers to the distribution of allelic effects, their interactions, and how segregating genetic variation contributes to differences in traits across individuals [7]. The genetic architectures of complex traits can be described by lists of trait-associated loci, their frequencies, and allele-specific effect sizes. Importantly, genetic architectures can vary across populations [7]. The relative importance of genetic and environmental effects varies by trait, and this can be quantified via estimates of SNP heritability (*h^2^_SNP_*) [8, 9]. An additional aspect of genetic architecture involves the extent to which trait-associated SNPs are in high linkage disequilibrium (LD) with surrounding regions of the genome. These differences may be particularly relevant to traits which are due to a small number of genes, since patterns of LD impact the ability of SNPs to tag causal variants and for GWAS findings to replicate across populations [10, 11]. There is also evidence that low LD SNPs have larger per-SNP heritabilities [12]. Furthermore, divergent evolutionary histories cause LD patterns to differ across populations [13]. The degree to which heritable variation is evenly distributed across the genome can differ depending on the trait [14, 15]. On one extreme are traits with Mendelian genetic architectures, like cystic fibrosis [16]. On the other extreme are highly polygenic traits like height [17]. Recently, an omnigenic model has been proposed - whereby traits have a set of core genes, but most of the heritability can be explained via indirect effects that are due to gene regulatory networks [18]. Consistent with the omnigenic model, causal variants for anthropometric and blood pressure traits have been found throughout the human genome [19]. Despite an awareness of the multiple ways that traits can differ, many aspects of genetic architecture have yet to be quantified in a comprehensive way.

GWAS findings can be used to generate polygenic scores (PGS), called polygenic risk scores in the context of hereditary diseases [20]. These scores enable traits to be predicted from genetic information, and they are commonly calculated by summing allele doses across all trait-associated loci and weighting by effect sizes [21–23]. Although the clinical utility of risk scores has received a significant amount of attention during the past few years, many PGS yield only modest case prediction accuracy and show weak correlations between predicted and actual trait values (R^2^ < 0.1) [24–26]. Even within the same ancestry, PGS accuracy can vary due to socio-economic status and genotype-environment interactions [27, 28]. These issues are even more pronounced when genetic predictions are applied to populations that have different ancestries than the original discovery population [29–31]. For example, predictions of anthropomorphic and blood-related traits generated from UK Biobank data perform better when applied to British individuals than Japanese individuals, while predictions generated from Biobank Japan data perform better when applied to Japanese individuals than British individuals [25]. In a landmark study of over 200 traits from the UK Biobank, Privé et al. found that the portability of genetic predictions is reduced in proportion to the genetic distance from the original discovery population [32]. Polygenic score accuracy also decays continuously over fine-scale differences in ancestry [33]. Furthermore, the predicted values of complex traits can vary between populations. These shifts in PGS distributions can either be due to ascertainment bias [34–36] or due to actual differences in traits [37, 38]. Although thousands of PGS have been generated to date [26], multiple knowledge gaps exist: Are there particular types of traits that are hard to predict from genetic data? Which traits have PGS that differ the most across populations?

Here, we leveraged GWAS effect sizes and polygenic score weights of 163 traits from the UK Biobank as well as population-specific LD scores and allele frequencies to quantify how different aspects of genetic architecture affect PGS performance. On top of SNP heritability, we developed two novel trait-specific metrics of genetic architecture (LD variability and genomic inequality) and identified relationships with three aspects of polygenic score performance: prediction accuracy in the source population, portability across populations, and trait divergence across populations, the latter two being novel trait-specific metrics as well. Finally, we identified suites of traits that have similar genetic architecture and PGS performance profiles. Notably, lifestyle and psychological traits were difficult to predict with genetic data, while also having PGS that generalize poorly across populations.

## Materials and methods

### PGS weights for 163 complex traits

Our paper builds upon the PGS weights previously generated by Privé et al. [32]. To our knowledge, this prior study contains the largest number of traits with multi-ancestry PGS performance metrics. After correcting for sex, age, deprivation index, and 16 principal components (PCs), Privé et al. used lasso penalized regression [39] to generate PGS from 391,124 British individuals of European descent from the UK Biobank (UKBB) [32]. We restricted our analyses to 163 traits that had publicly available PGS weights, SNP heritability, and accuracy statistics, and an equivalent GWAS computed in the Pan UK Biobank (described below). These traits were grouped into four categories: 53 biological measures (including blood phenotypes), 48 diseases, 24 lifestyle/psychological traits, and 38 physical measures (such as height and weight). We only included binary traits with a prevalence above 1% in the UK Biobank due to concerns about statistical power and reliability of trait metrics. A total of 67 binary and 96 quantitative traits were analyzed here. Due to the diminished statistical power inherent to binary trait GWAS, the bulk of our analyses focused on quantitative traits [40]. A full list of traits, as well as summary statistics of genetic architecture and PGS performance, can be found in Table S1.

### GWAS Effect Sizes

Since PGS weights were calculated using lasso, which can be considered to use an unjustified Laplace prior [41], their posterior distribution may not be representative of true effect sizes, especially after LD pruning. To obtain less biased SNP weights, we extracted trait-specific effect sizes from the Pan UK Biobank (Pan UKBB, https://pan.ukbb.broadinstitute.org), a repository of ancestry-specific GWAS, heritability, and LD score results for over 500,000 individuals in the UK Biobank. The Pan UKBB split individuals into six ancestries: African (AFR), Admixed American (AMR), Central/South Asian (CSA), East Asian (EAS), European (EUR), and Middle Eastern (MID), the latter of which was excluded from our analyses due to a lack of equivalent samples in the 1000 Genomes Project phase 3 data [42]. Note that each ancestry category is heterogeneous – genetic ancestries exist along a continuum [33, 43]. GWAS results were filtered to include only variants with allele frequency and effect size data for all ancestries and pruned using Plink [44] (p-value < 10^-5^, r^2^ < 0.2), and corrected for the Winner’s Curse [45]. When allele frequency data was missing from the Pan UKBB, we used continental allele frequencies from the 1000 Genomes Project [44]. We then matched traits in the Pan UKBB to trait fields in Privé et al.’s analyses (Table S1). Because some UKBB categorical variables were transformed differently by Privé et al. and the Pan UKBB, for a trait to be considered quantitative in our analyses, we required that both sources of data listed the trait as quantitative.

### Genetic variance contributions

We leveraged effect size and allele frequency information to identify the most important trait-associated SNPs. Specifically, alleles with large effect sizes and minor allele frequencies close to 50% contribute more to the heritable variation of a trait than alleles with small effect sizes and minor allele frequencies close to zero. As described in earlier work [46, 47], the contribution of a SNP to the total genetic variance of a trait under Hardy-Weinberg equilibrium and an additive polygenic model is given by:

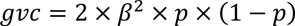

where *gvc* refers to the genetic variance contribution, *β* is effect size per allele copy obtained from the Pan UKBB GWAS meta-analysis, and *p* is the frequency of the reference allele for a given ancestry. *gvc* values were computed for each Pan UKBB ancestry.

### Quantifying SNP heritability, LD variability, and genomic inequality

SNP heritability (*h^2^_SNP_*) estimates for each trait were previously calculated by Privé et al. [32] using LDpred2-auto [48].

LD scores from the Pan UKBB were used to derive a metric of how much the SNPs associated with a given trait have LD patterns that vary by ancestry. Ancestry-specific LD scores were previously computed using LD score regression [49]. Note that SNPs with low LD scores tag fewer other SNPs. For each combination of trait and ancestry, we calculated the *gvc*-weighted arithmetic mean LD score of the top 500 trait-associated SNPs within each ancestry. The standard deviation (𝜎*_LDscore_*) and mean (𝜇*_LDscore_*) across ancestries were then found for each trait. We define LD variability as the coefficient of variation of ancestry-specific LD scores for each trait:

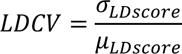

Continental ancestries from the Pan UKBB (AFR, AMR, CSA, EAS, and EUR) were used in the above calculations.

We quantified the inequality in the genetic variance contribution of top trait-associated SNPs using a novel application of Gini coefficients. These coefficients have typically been used in economics to calculate wealth or income inequality [50], and they range between zero (maximum equality) and one (maximum inequality). Here, we used Gini coefficients to quantify the extent that *gvc* is evenly distributed among the top 500 SNPs by *gvc*. Focusing on the top 500 SNPs maximizes the dynamic range of Gini coefficients and the ability to capture a significant amount of genetic variance (Fig. S1). Regardless, the number of top SNPs chosen does not substantially affect the ranking of traits by their Gini coefficient (Fig. S2). For traits that had fewer than 500 independently significant SNPs, dummy SNPs with *gvc* values of zero were included so that Gini coefficient calculations always consisted of exactly 500 elements. Gini coefficients were calculated using the following equation, which requires that SNPs be sorted by *gvc* in ascending order:

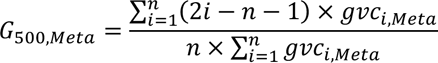

where *n* = 500 and *gvc_i_* is the summed genetic variance contribution of the *i*^th^ SNP using allele frequencies from the Pan UKBB meta-analysis. Using https://github.com/oliviaguest/gini as a guide, we implemented a computationally efficient R script that uses the above equation to compute Gini coefficients for each trait. Ancestry-specific Gini coefficients were also computed from Pan UKBB data.

### Quantifying PGS accuracy, portability, and divergence

Here, PGS accuracy refers to how well genetic predictions work when individuals are ancestry-matched to the original training set (i.e., British individuals from the UK). For each trait, partial correlations between predicted and actual trait values for Privé et al.’s UK ancestry group (ρ*_UK_*) were used to quantify PGS accuracy [32]. These partial correlations were generated using the residuals of actual trait values and PGS after correcting for the following covariates: age, sex, birth date, deprivation index, and population structure (16 PCs) [32]. This statistic was calculated using genome-wide PGS weights.

Partial correlations between predicted and actual trait values were obtained for eight ancestries in Privé et al.’s data [32]: UK, Poland, Italy, Iran, India, China, Caribbean, and Nigeria. Using this information, we derived a portability index that quantifies how PGS accuracy diminishes with increased genetic distance from the original training population (UK). For each trait and ancestry group, PGS accuracy relative to the UK ancestry group was found by dividing the partial correlation between predicted and actual trait values for each ancestry group by the partial correlation for the UK ancestry group. Similarly, as per Privé et al. [32], the geometric mean position of each ancestry group in 16-dimensional PC space was used to obtain the Euclidean genetic distance between each ancestry group and the UK ancestry. For each trait, relative PGS accuracy was plotted against genetic distance to the UK ancestry group, and linear regression was used to quantify PGS portability. We define the slope of the regression line for each trait (*m*) as the portability index of that trait. Regression lines were required to pass through the UK datapoint (0,1). Noisy PGS accuracy statistics can cause some traits to have slopes above 0, which we corrected by manually setting *m* to be 0, i.e., perfect portability.

We also developed a summary statistic that quantifies how much PGS distributions have diverged across populations. As mentioned above, we recreated eight of the ancestry groups found in the UKBB using the procedure described in Note A of Privé et al. [32]. Numbers of individuals for each ancestry group in the UKBB were downsampled to 1,234, the smallest number of samples in any one ancestry group. PGS scores were calculated using the –score command in Plink 2 [51], which sums allele dose (*d_j,k_*) multiplied by effect size (*B_k_*) across all *L* trait-associated SNPs for each trait:

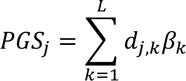

where *j* indexes each individual and *k* indexes each trait-associated SNP. We then generated PGS distributions for each ancestry group and all 163 traits. Because PGS are calculated by summing the effects of multiple independent SNPs, these distributions tend to be normally distributed. We then log-transformed the F-statistic from a one-way ANOVA to derive a metric (*D*) which quantifies population-level shifts in PGS distributions for each trait:

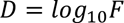

where *F* refers to the ratio of between-ancestry group variability to within-ancestry group variability. We note that meaningful comparisons of *D* statistics for different traits require that same number of individuals from each ancestry group were analyzed for each trait, as was the case in our study design.

### Comparisons between different aspects of complex traits

Our manuscript focuses on three summary statistics of genetic architecture (*h^2^_SNP_*, *LDCV*, *G_500,Meta_*) and three summary statistics of PGS performance (ρ*_UK_*, *m*, and *D*) for each trait. To compare different aspects of genetic architecture and PGS performance, we obtained a linear best fit for each pair of summary statistics, generating correlation coefficients and p-values. Because 15 pairwise comparisons were made, a false discovery rate (FDR) [52] adjustment was applied to each p-value using the p.adjust() command in R. Note that the Benjamini-Hochberg FDR procedure can cause p-values to clump.

We also assessed how distributions of six summary statistics (*h^2^_SNP_*, *LDCV*, *G_500,Meta_*, ρ*_UK_*, *m*, and *D*) vary for different types of traits. First, we compared the summary statistic distributions of binary traits with the distributions of quantitative traits. We then compared the summary statistic distributions of quantitative lifestyle/psychological traits with the distributions of other quantitative traits. Wilcoxon rank sum tests [53] were used for these summary statistic comparisons, and FDR-adjusted p-values were used to correct for multiple comparisons. Principal component analysis (PCA) was also used to identify quantitative traits with similar summary statistics. Specifically, we applied the prcomp() function in R on a 96 by 6 array containing genetic architecture and PGS performance summary statistics for all quantitative traits. PCA plots used a modified version of the *ggbiplot* package in R, with 68% probability ellipses (+/- one standard deviation) shown for different groups of traits.

## Results

### Different metrics of genetic architecture and polygenic score performance

Here we describe six trait-specific metrics, each of which captures a different aspect of genetic architecture or PGS performance. Two of these metrics were previously calculated (SNP heritability and PGS accuracy), while four are novel (LD variability, genomic inequality, portability, and divergence). Examples of traits with low and high values of these novel metrics are shown in Fig. 1. A full list of trait-specific metrics can be found in Table S1.

**Fig. 1.**
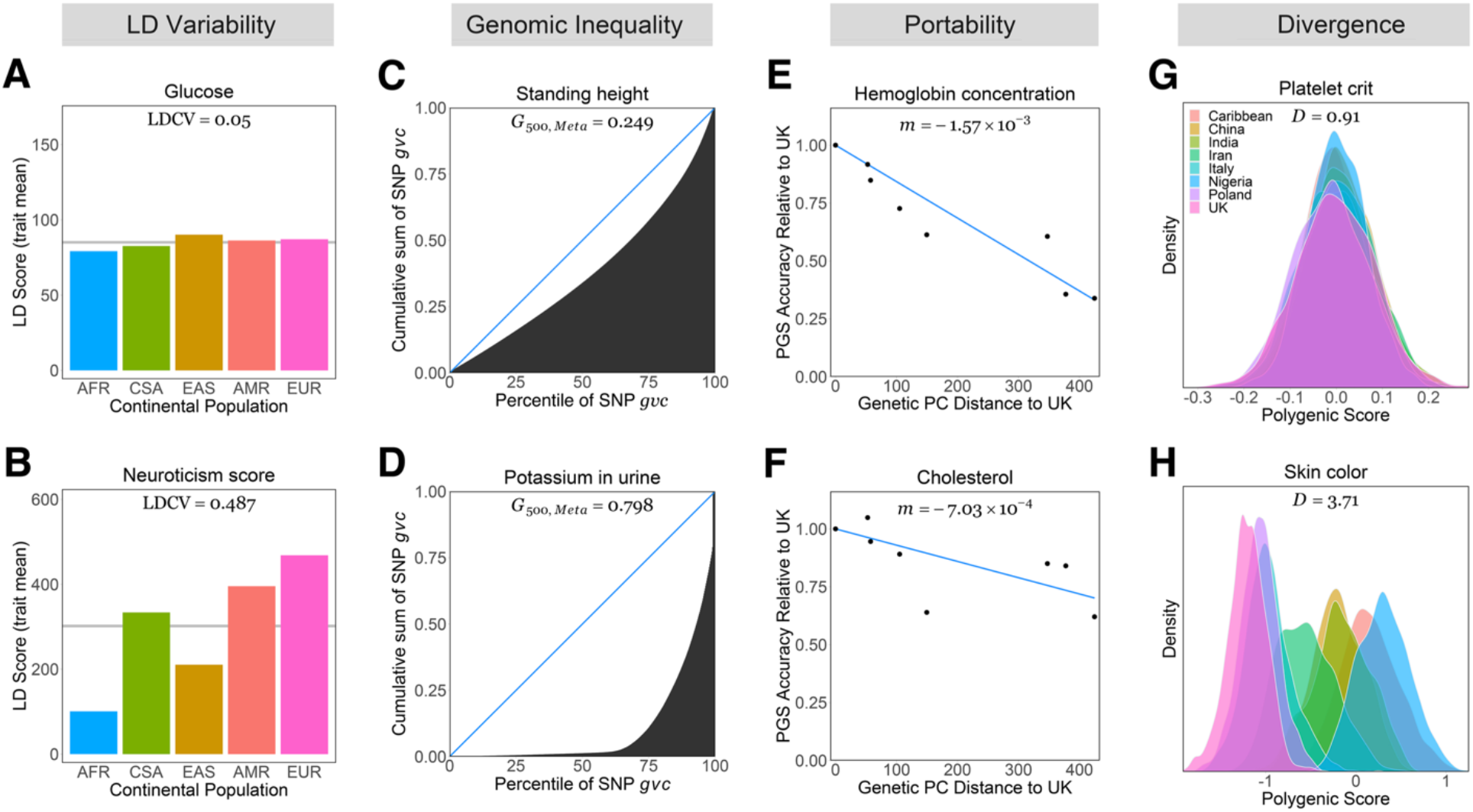
Examples of trait-level differences in LD variability, portability, and divergence. LD variability (*LDCV*) quantify how much LD scores vary across ancestries. Glucose level (**A**) has low LD variability while neuroticism (**B**) has high LD variability. Gini coefficients (*G_500,Meta_*) quantify how unequally the genetic variance of each trait is distributed across the top trait-associated SNPs. Lorenz curves of the top 500 independent SNPs are shown here. Height (**C**) has a more equal distribution of genetic variance while potassium concentration (**D**) in urine has a more unequal distribution. Portability statistics (*m*) quantify how well genetic predictions generalize to different ancestry groups. Predictions of hemoglobin concentration (**E**) have low portability while predictions of cholesterol levels (**F**) have high portability. Divergence statistics (*D*) quantify the extent that PGS distribution means differ across ancestry groups. Ancestry-specific PGS distributions for platelet crit (**G**) have a high amount of overlap while PGS distributions for skin color (**H**) are highly divergent.

We used existing metrics computed by Privé et al. [32] for SNP heritability (*h^2^_SNP_*) and PGS accuracy (ρ_UK_). SNP heritability quantifies the proportion of the total phenotypic variance of a trait that can be explained by common SNPs, and PGS accuracy measures the correlation between actual and predicted values of traits. Quantitative traits with high values of both SNP heritability and PGS accuracy include standing height (*h^2^_SNP_* = 0.546, ρ_UK_ = 0.634), mean thrombocyte volume (*h^2^_SNP_* = 0.392, ρ_UK_ = 0.603) and total bilirubin (*h^2^_SNP_* = 0.361, ρ_UK_ = 0.605). By contrast, urine content of microalbumin (*h^2^_SNP_* = 0.0365) and potassium (*h^2^_SNP_* = 0.0435) were the two quantitative traits with the lowest SNP heritability, while general happiness with life (ρ_UK_ = 0.0640) and with health (ρ_UK_ = 0.0697) were the two quantitative traits with the lowest PGS prediction accuracy.

Our LD variability statistics measure how different the LD patterns of trait-associated SNPs are across ancestries. Glucose level (Fig. 1A) was the quantitative trait whose top SNPs differed the least in LD scores across ancestries (*LDCV* = 0.050), meaning that associated SNPs’ ability to tag causal variants are more likely to be preserved across populations. Other low LD variability traits include cystatin C levels (*LDCV* = 0.058) and pulse rate (*LDCV* = 0.074). Traits with high LD variability included lifestyle-related traits like water intake (*LDCV* = 0.534), general happiness with health (*LDCV* = 0.533), and neuroticism (*LDCV* = 0.489, Fig. 1B), as well as urine content concentrations of potassium (*LDCV* = 0.532) and microalbumin (*LDCV* = 0.0526).

We also used a novel application of Gini coefficients to quantify the genomic inequality of complex traits. A low Gini coefficient indicates that the genetic variance of a trait tends to be equally distributed among top associated SNPs, i.e., a more polygenic architecture. By contrast, a high Gini coefficient indicates that a large portion of a trait’s genetic variance is explained by a small set of SNPs, i.e., a more Mendelian genetic architecture. We emphasize that the Gini coefficient is not a measure of how many SNPs are associated with a trait, but instead how unequally the *gvc* of the top trait associated SNPs is distributed. Quantitative traits with the lowest and highest Gini coefficients are shown in Table 1. Lorenz curves can be used to visualize how Gini coefficients summarize differences in *gvc* distributions for complex traits. For example, the *gvc* of the top SNPs associated with standing height are relatively evenly distributed (*G*_500,Meta_ = 0.249, Fig. 1C). An example of a high Gini trait is potassium concentration in urine (*G*_500,Meta_ = 0.798, Fig. 1D). We note that traits with large Gini coefficients also tend to have a larger proportion of their summed *gvc* in the top 500 SNPs (Fig. S1). In addition, the rank orders of Gini coefficients were largely robust to the number of top SNPs that were examined (Fig. S2). Using continental ancestry-specific allele frequencies in the calculation of *gvc*, we obtained ancestry-specific Gini coefficients (Fig. S3). Non-African Gini coefficients (Fig. S3A-D, r = 0.820 to 0.889) had higher correlations with meta-analysis Gini coefficients than what was observed for African Gini coefficients (Fig. S3E, r = 0.661). These differences are likely due to the large fraction of Eurasian individuals in the Pan UKBB meta-analysis. We posit that large effect non-European variants may cause some quantitative traits to have higher *G*_500,Meta_ than *G*_500,EUR_ (Fig. S3A). Conversely, many quantitative traits have a higher *G*_500,AFR_ than *G*_500,Meta_ (Fig. 3SE). This pattern is likely due to ascertainment bias, i.e., trait-associated SNPs in the (largely European) meta-analysis are not guaranteed to polymorphic in African individuals.

**Table 1.** Quantitative traits with the lowest and highest genomic inequality statistics (Gini). Higher values of G_500,Meta_ are indicative of traits with more Mendelian genetic architectures.

| Rank | Trait | $G_{500, \text{Meta}}$ |
| --- | --- | --- |
| 1 | First sex age | 0.187 |
| 2 | Systolic blood pressure | 0.220 |
| 3 | Forced expiratory volume | 0.221 |
| 4 | Hip circumference | 0.225 |
| 5 | Diastolic blood pressure | 0.228 |
| ... | ... | ... |
| 92 | General happiness | 0.943 |
| 93 | Menstrual cycle length | 0.944 |
| 94 | General happiness in own health | 0.958 |
| 95 | Carotid intima-media thickness | 0.963 |
| 96 | Electrocardiogram P duration | 0.963 |

Another aspect of PGS performance is the portability of results across different populations. For each trait, we plotted the relative PGS accuracy for different ancestry groups, applied a linear model, and used the slope (*m*) to quantify the portability of genetic predictions. If *m* = 0, genetic predictions work equally well for each ancestry group. By contrast, strongly negative slopes (*m* < 0) indicate increasingly poor predictive power relative to the UK ancestry group. Here, we use two examples to illustrate how genetic predictions of complex traits can differ in their portability. Hemoglobin concentration has a low portability statistic (*m* = −0.00157, Fig. 1E). By contrast, cholesterol has a high portability statistic (*m* = −0.000703, Fig. 1F).

We also developed a novel metric of PGS divergence across populations (*D*). This metric was calculated by examining the PGS distributions for eight different ancestries use PGS weights from Privé et al.’s [32] and converting F statistics from a one-way ANOVA to a log_10_ scale. *D* statistics near zero arise when ancestry-specific PGS distributions have similar means; higher values of *D* statistics arise when ancestry-specific PGS distributions have different means. Because of this, *D* statistics can be used to quantify ancestry-specific shifts in PGS distributions. Lists of the most and least divergent traits are shown in Table 2 and Table S1. An example of a trait with minimal PGS divergence is benign platelet crit (*D* = 0.91, Fig. 1G). An example of a trait with substantial divergence between ancestries is skin color (*D* = 3.71, Fig. 1H).

**Table 2.**
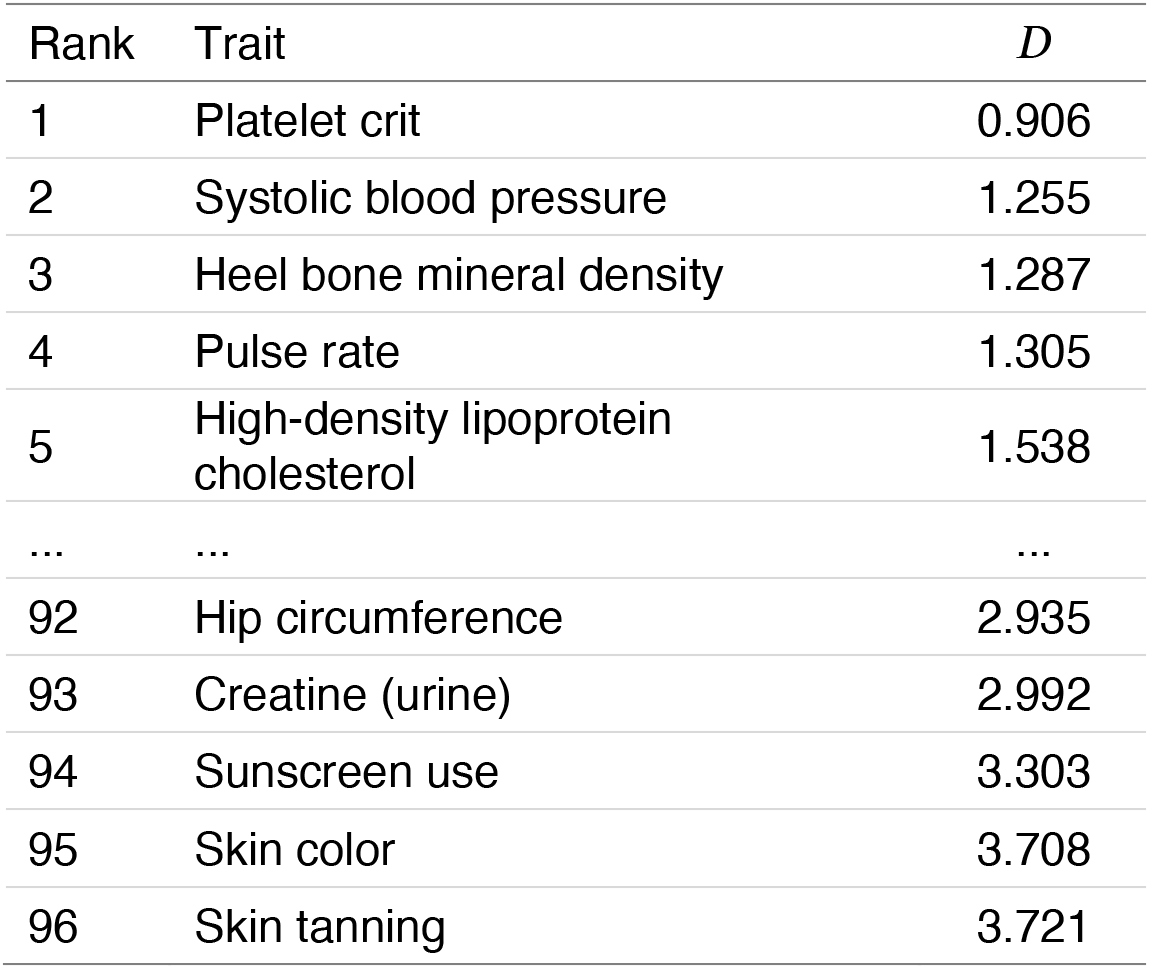
Quantitative traits with the lowest and highest PGS divergence statistics. Higher values of *D* are indicative of traits that have PGS distribution means that differ more across ancestries.

### Summary statistics differ for binary and quantitative traits

One natural way to classify traits is whether they are binary or quantitative (the latter including both ordinal and continuous traits). All the disease traits analyzed in our study are binary. In addition, the statistical power to detect genetic associations differs for binary and quantitative traits [40]. Both of these details can affect estimates of genetic architecture and PGS performance. Indeed, we observed differences in the summary statistics of binary and quantitative traits (Fig. 2). On average, binary traits tended to have a lower SNP heritability (p = 6.03 x 10^-9^) than quantitative traits, but higher LD variability (p = 4.00 x 10^-9^) and Gini (p = 8.52 x 10^-17^, Wilcoxon rank sum tests for all comparisons). The latter is most certainly driven by the decreased power in GWAS of binary traits [37] resulting in fewer overall SNP associations being found. Focusing on different aspects of PGS performance, we found that PGS accuracy was much lower for binary traits than quantitative traits (p = 3.41 x 10^-21^). For example, partial correlations between predicted and actual trait values were lower for hypertension (ρ_UK_ = 0.188) than for systolic blood pressure (ρ_UK_ = 0.255). Binary traits also had slightly lower (p = 0.006) and a wider range of portability statistics than quantitative traits (Fig. 2). However, we note that our portability statistic is less reliable at lower PGS accuracies. Finally, we note that divergence statistics were similar for binary and quantitative traits (p = 0.295). Due to low SNP heritabilities and PGS accuracies of binary traits, our subsequent analyses focus on quantitative traits.

**Fig 2.**
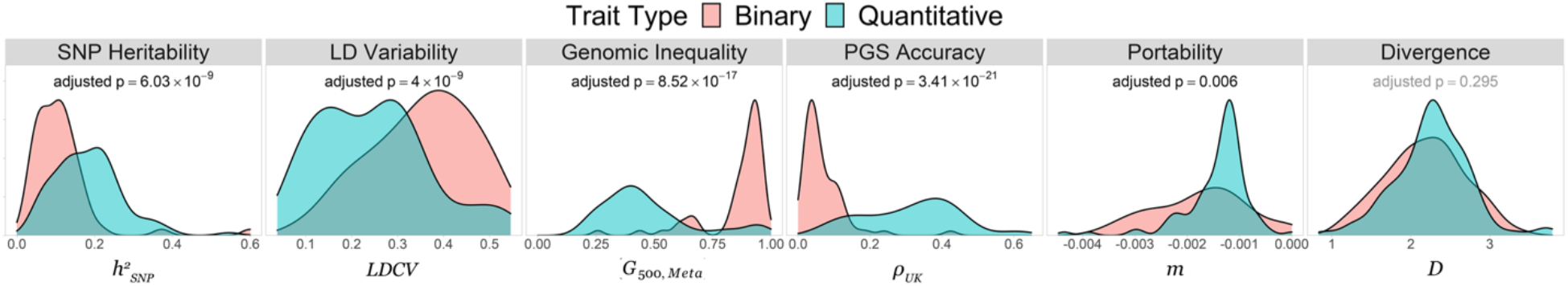
Distributions of genetic architecture and polygenic score performance metrics for binary and quantitative traits. All p-values are obtained from Wilcoxon rank sum tests, and statistically significant (FDR-adjusted p < 0.05) results are shown in bold.

### Comparisons between different aspects of genetic architecture

SNP heritabilities, LD variability, and Gini coefficients capture different aspects of genetic architecture (upper left part of Fig. 3). High LD variability was associated with neither SNP heritability (r = −0.083, p = 0.453) nor Gini coefficients (r = 0.131, p = 0.298). However, SNP heritability was negatively correlated with Gini (r = −0.444, p = 2.87 x 10^-5^), meaning that less heritable traits tended to have less equally distributed genetic variance among top SNPs. We note that there was a lack of quantitative traits that had both a high SNP heritability and a high Gini coefficient. By contrast, there were many traits with both a low SNP heritability and a low Gini coefficient, particularly lifestyle-related traits such as household income (*h^2^_SNP_* = 0.0979, G_500,Meta_ = 0.273), overall health rating (*h^2^_SNP_* = 0.104, G_500,Meta_ = 0.274), and chronotype (*h^2^_SNP_* = 0.127, G_500,Meta_ = 0.233). Traits exhibiting such patterns appear to have a truly polygenic architecture, as their low SNP heritability is not simply a result of having fewer trait-associated loci.

**Fig. 3.**
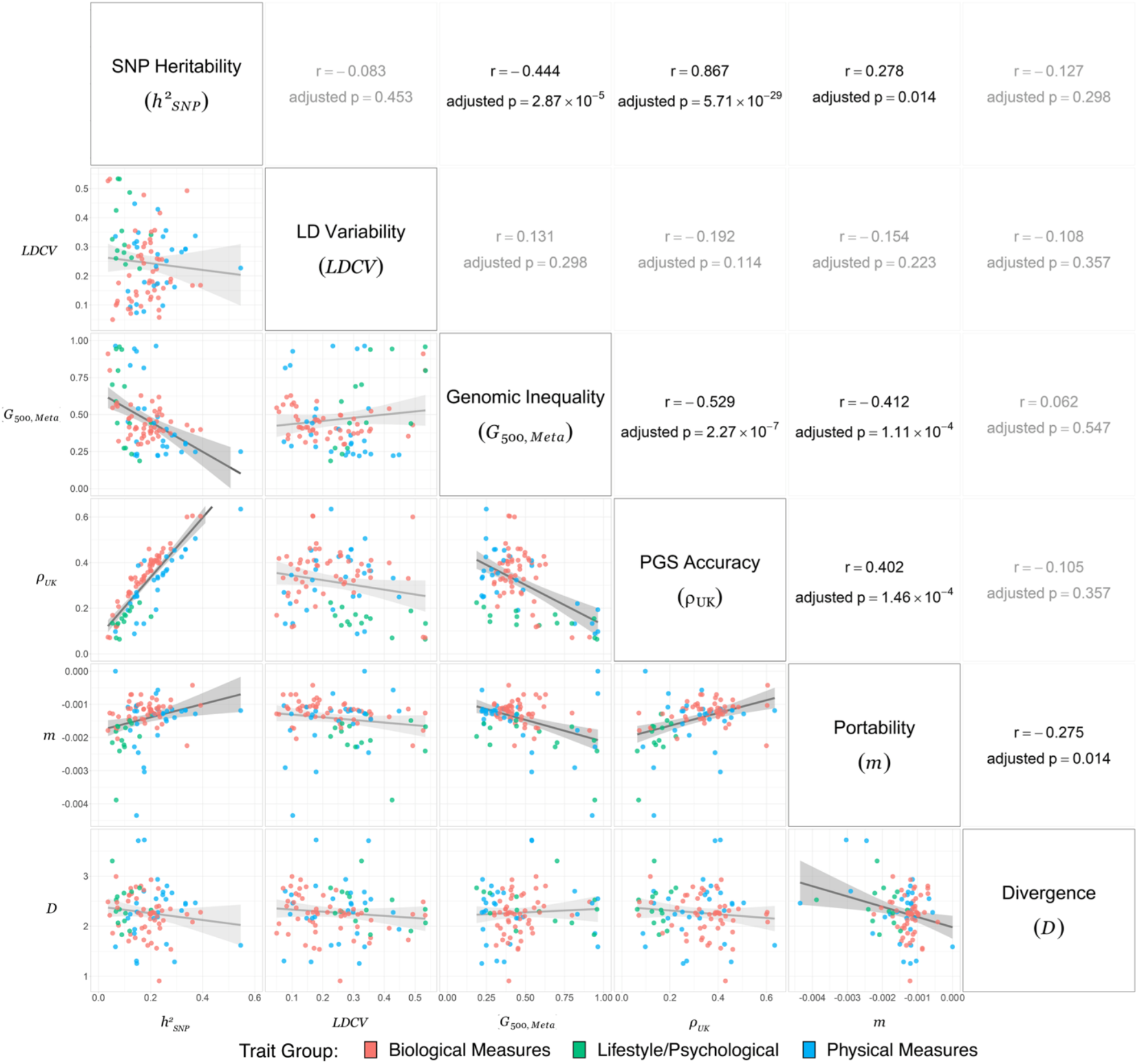
Comparisons between different aspects of genetic architecture and PGS performance for 96 quantitative traits. Each scatterplot corresponds to a different pair of summary statistics., each data point corresponds to a different quantitative trait, and colors are indicative of trait group. All p-values are FDR-adjusted, and statistically significant results are shown in bold. Summary statistics for all traits can be found in Table S1.

### Comparisons between different aspects of polygenic score performance

Complex traits vary in multiple aspects of PGS performance: accuracy, portability, and divergence (lower right part of Fig. 3). We found that PGS accuracy and portability were positively correlated (r = 0.402, p = 1.46 x 10^-4^), meaning that traits which are easier to predict in the UK also tend to have a more modest falloff in prediction accuracy with genetic distance to the source population. An example of a trait with high PGS accuracy and high portability is mean platelet volume (ρ*_UK_* = 0.603, *m* = −0.00103). An example of a trait with low PGS accuracy and low portability is general happiness (ρ*_UK_* = 0.0697, *m* = −0.00388). Portability statistics are noisier for traits that are hard to predict in British individuals from the UK. Because of this, values of *m* are highly variable when ρ*_UK_* is close to zero. We note that divergence and portability statistics are negatively correlated (r = −0.275, p = 0.014). This pattern can arise from a combination of ascertainment bias and allele frequency differences between populations [54]. There were no broad trends when trait divergence was compared to PGS accuracy (r = −0.105, p = 0.357).

### Relevance of genetic architecture to polygenic score performance

How do SNP heritabilities, LD variability, and genomic inequality relate to PGS accuracy, portability, and divergence? Although not all combinations of genetic architecture and PGS performance yielded clear associations, a few notable patterns can be seen in the top right part of Fig. 3. As expected, traits with a high SNP heritability tended to have a high PGS accuracy (r = 0.867, p = 5.71 x 10^-29^). Interestingly, every lifestyle trait’s PGS accuracy underperformed with respect to their heritability as predicted by the trend line (mean residual = −0.0581, two-sided Wilcoxon rank sum test: p = 1.22 x 10^-4^). Physical measures’ PGS accuracies also significantly underperformed with respect to the trendline (mean residual = −0.0463, p = 1.56 x 10^-4^) while biological measures overperformed (mean residual = 0.0407, p = 1.76 x 10^-7^). We note that heritability refers to the proportion of phenotypic variance that is due to genetic effects in a single population (i.e., it is a population-specific concept). This suggests that *h^2^_SNP_* estimates may not be that informative about how well predictions generalize across populations. Indeed, despite being positively associated with portability (r = 0.278, p = 0.014), SNP heritabilities were less informative about the portability of polygenic predictions than PGS accuracy in the source population (t-test on a Fisher’s r to z transform: p = 0.012) and were non-informative about the divergence of predicted trait values (r = −0.127, p = 0.298). Traits with unequal *gvc* distributions (high Gini) tended to have less accurate PGS predictions (r = −0.529, p = 2.27 x 10^-7^), a pattern that was likely driven by low SNP heritability traits. Furthermore, high Gini traits also tended to have less portable PGS predictions (r = −0.412, p = 1.11 x 10^-4^), perhaps due to concentrated genetic effects being more prone to noise across populations due to allele frequency differences. Notably, LD variability was not significantly associated with any other metric. Particularly surprising was the absence of a significant relationship between LD variability and portability (r = −0.154, p = 0.223), as LD variability is a metric that quantifies differential tagging of GWAS variants across ancestries.

### Limitations of polygenic scores for lifestyle and psychological traits

A PCA plot generated from six summary statistics of genetic architecture and PGS performance further demonstrates that lifestyle and psychological traits have different profiles than other quantitative traits (Fig. 4A). Arrows in this plot indicate regions of PCA space that are associated with higher values of each summary statistic, recapitulating our earlier findings: higher heritability is strongly associated with higher PGS accuracy and higher portability. We also note that the arrows for LD variability and divergence point in different directions. Lifestyle and psychological traits form a noticeable cluster in the region of PCA space pertaining to lower heritability, prediction, and portability statistics. By contrast, biological measures and physical measures occupy overlapping regions of PCA space, which reflects that these two trait groups are more varied in their genetic architecture and PGS performance. We also point out that despite being centered in same region, biological measures (i.e., endophenotypes) were more closely clustered than physical measures.

**Fig. 4.**
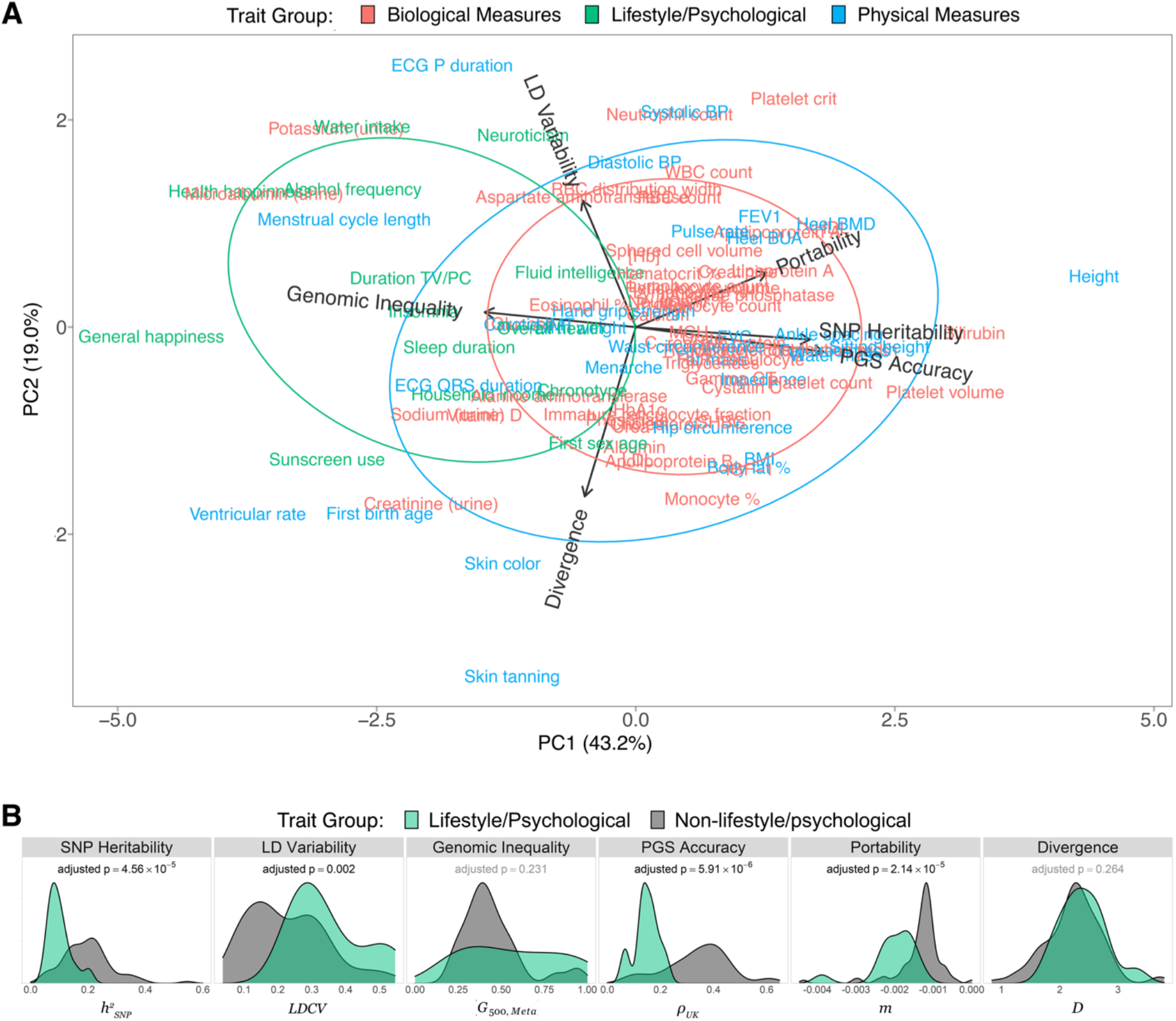
Lifestyle and psychological traits have different genetic profiles than other quantitative traits. **A** PCA plot generated from six summary statistics of genetic architecture and PGS performance (*h^2^_SNP_*, *LDCV*, *G_500,Meta_*, ρ*_K_*, *m*, and *D*). Arrows indicate higher values of each summary statistic in PCA space, and each summary statistics has been scaled to have unit variance. All traits shown here are quantitative, and trait names are colored by trait group. Ellipses indicate 68% (+/- one standard deviation) confidence intervals. **B** Distributions of genetic architecture and polygenic score performance metrics for quantitative lifestyle and psychological traits vs. other quantitative traits. All p-values are obtained from Wilcoxon rank sum tests, and statistically significant (FDR-adjusted p < 0.05) results are shown in bold.

Given the distinct cluster of lifestyle traits, we compared their summary statistics to those of the remaining two quantitative trait groups. Examples of these complex behavioral traits include alcohol frequency, chronotype, happiness, neuroticism, and water intake. Overall, there are noticeable differences in the genetic architecture and PGS performance of different types of quantitative traits (Fig. 4B). On average, lifestyle and psychological traits have lower SNP heritabilities than other quantitative traits (p = 4.56 x 10^-5^). This pattern is consistent with the importance of environmental effects for behavioral traits. Low heritability also has a knock-on effect of reducing the effectiveness of polygenic predictions. Indeed, we found that PGS accuracy was much lower for lifestyle and psychological traits (p = 5.91 x 10^-6^). Furthermore, we note that there was a clear lack of lifestyle and psychological traits with high ρ*_UK_* statistics (Fig. 4B). Genetic predictions of lifestyle and psychological traits were also less portable than other quantitative traits (p = 2.14 x 10^-5^) and slightly more variable in their LD scores (p = 0.002). No significant differences between the means of lifestyle/psychological traits and other quantitative traits were present for Gini coefficients (p = 0.231) and PGS divergence (p = 0.264). However, we do note that lifestyle and psychological traits had much greater variance in their Gini coefficients, consisting of both high and low G_500,Meta_ values, while other quantitative traits mostly occupied a narrow band near the middle of the overall distribution. Taken together, Fig. 4B reveals that genetic predictions of lifestyle and psychological traits are severely limited given our present knowledge of the genetic basis of these traits.

## Discussion

Overall, we found that complex traits have a broad range of genetic architectures, some of which contribute to differences in PGS performance. Our results indicate that highly heritable traits are easier to predict when individuals are ancestry-matched to the original GWAS cohort – a finding that is consistent with expectations from statistical genetics [55]. However, SNP heritability was somewhat less informative when it came to the portability of genetic predictions. Intriguingly, LD variability was largely uninformative regarding the portability of polygenic predictions. This suggests that factors other than the differential tagging of causal SNPs drive the reduction in polygenic score accuracy across populations. Furthermore, since the meta-analysis consisted almost entirely of European samples, we caution that LD variability as measured here is hindered by ascertainment bias, and thus it may not fully capture true LD differences across populations.

Shifts in PGS distributions are due to allele frequency differences between populations. Because of this, one might expect to find less overlap in the PGS distributions of high Gini traits than for low Gini traits. The reasoning here is that allele frequency differences at different SNPs can average out if traits are highly polygenic, but greater noise between populations is possible if traits have more Mendelian architectures. However, our summary statistics of trait divergence were largely independent of genomic inequality. This suggests that other phenomena like natural selection [56] and ascertainment bias [34] are drivers of ancestry-specific shifts in PGS distributions. Indeed, hair and skin color, which are among the most divergent traits in our study, have previously been implicated in scans of selection [57]. We also note that natural selection can erode the portability of polygenic predictions [58, 59].

Our analyses also shed light on the omnigenic model. Gini coefficients varied substantially across traits, a pattern that was not an artifact of focusing on the top 500 independent SNPs for each trait (Fig. S2). Our results demonstrated that genetic architectures are often trait-specific, and that core genes can potentially make outsized contributions to the overall genetic variance of a trait, as many traits had a high Gini coefficient. That said, the omnigenic model proposes that genetic effects cascade through cellular regulatory networks, as expression of core genes ends up affecting gene expression at other genes [18]. This might also explain why transcriptional risk scores can potentially outperform genetic risk scores [60].

When large well-powered GWAS are conducted on diverse cohorts, rare family-specific or ancestry-specific variants are more likely to identified. However, genetic associations involving these private alleles are unlikely to yield portable predictions. Indeed, a height GWAS of 5.4 million individuals found that SNP heritability clusters in genomes, and that out-of-sample prediction accuracy was lower for individuals who did not have European ancestry [61]. There is also evidence that pruning sets of trait-associated SNPs can lead to improved PGS performance among diverse populations [62].

We mention that PGS are not immune to controversy – especially when it comes to lifestyle and psychological traits. Some have envisioned a world where PGS for educational attainment might be used inform the allocation of resources to those who have the most need [63]. This has spurred intense debate about both efficacy of polygenic predictions for behavioral traits and whether they should be used in a public policy setting [64, 65]. Others have gone a step further and advocated using PGS to screen embryos for cognitive traits [66], a position that has received well-warranted criticism [67, 68]. Regardless of the specific trait, there are major challenges to polygenic screening of embryos [69–71]. Polygenic predictions of complex behavioral traits are particularly problematic. As seen in Fig. 4B, lifestyle and psychological traits are difficult to predict, which means that any downstream applications of PGS for these traits would be deeply flawed. This issue is particularly acute when PGS are applied to populations that have ancestries that differ from the original GWAS population, given their low portability. Ultimately, genetic predictions of traits like alcohol intake, general happiness, income, or educational attainment in non-European populations should be treated with extreme skepticism; racist claims about the supposed intellectual superiority of any particular ancestry are genetically untenable.

Lastly, we address some limitations of our study. Genetic architecture and PGS performance comparisons across populations are less reliable when sample sizes vary by population. Because of this, we strongly advocate for the continuation of initiatives to recruit diverse biobank cohorts, such as in the All of Us program [72] and Our Future Health (https://ourfuturehealth.org.uk). Furthermore, the UK Biobank is not fully representative of the UK population, as participants tend to be older, healthier, and wealthier than the general population [73]. Likewise, initiatives to recruit participants from low socio-economic standing is paramount for achieving equitable benefits of genomic medicine. Additionally, GWAS approaches are susceptible to model misspecification [74], and thus any GWAS-based method of quantifying genetic architecture may not capture the true genetic architecture of each trait. We attempted to circumvent this constraint by correcting for the Winner’s Curse and by focusing on SNPs whose trait associations we were most confident about. We concede that this required that our implementation of the Gini coefficient focus on the top trait-associated SNPs, as opposed to the entire genome. As previously mentioned, our portability metric is noisy at low PGS accuracies, and consolidating individuals into discrete ancestral groups to compute portability may simplify the continuous nature of PGS accuracy decay along genetic principal components. Finally, although our PGS divergence statistic successfully captured expected differences in genetic trait liability across ancestries (such as skin color), this metric is also biased towards genetic liability captured in European SNPs and thus may not fully capture true PGS divergence.

## Conclusion

We note that the summary statistics examined here are not exhaustive. Going forward, future studies will be able to explore additional aspects of genetic architecture and PGS performance. For example, some traits are highly canalized, while others show evidence of substantial PGS-by-environment interactions [75]. Epistatic interactions also contribute to the genetic architecture of complex traits, and this information can be incorporated into predictive models [76]. Future studies are also likely to benefit from the development of portability metrics that can account for both continuous genetic ancestry as well as environment. Finally, we note that PGS generated from multi-ancestry cohorts are more likely to yield portable predictions [77]. Nevertheless, we still expect there to be significant limitations to the genetic prediction of complex behavioral traits, especially across populations.

## List of abbreviations

AFR: African ancestry
AMR: Admixed American ancestry
CSA: Central/South Asian ancestry
EAS: East Asian ancestry
EUR: European ancestry
FDR: false discovery rate
*gvc*: genetic variance contribution
GWAS: genome-wide association study
*h^2^_SNP_*: SNP heritability
LD: linkage disequilibrium
*LDCV*: LD score coefficient of variation (a measure of LD variability)
*Meta*: Pan UK Biobank meta-analysis
MID: Middle Eastern ancestry
PCA: principal component analysis
PGS: polygenic score
SNP: single nucleotide polymorphism
UKBB: UK Biobank

## Acknowledgements

We thank study participants from the UK Biobank. In addition, we thank Greg Gibson, Sini Nagpal, King Jordan, Aaron Pfennig, Mimi Holness, and other members of the Center for Integrative Genomics at Georgia Institute of Technology for their helpful suggestions and feedback.

## Authors’ contributions

N.C.: methodology, formal analysis, data curation, visualization, and writing; A.H.: methodology, formal analysis, data curation, visualization, and writing; J.L.: conceptualization, funding acquisition, methodology, supervision, visualization, and writing.

## Availability of data and materials

Datasets supporting the conclusions of this article are available in multiple repositories.

• UK Biobank data can be requested via: https://www.ukbiobank.ac.uk/enable-your-research.

• PGS weights and ancestry-specific partial correlations (PGS accuracy statistics) are described in Privé et al. [32], and they can be accessed at: https://figshare.com/articles/dataset/Effect_sizes_for_215_polygenic_scores/14074760/2

• SNP heritabilities described in Privé et al. [32], and they can be accessed at: https://github.com/privefl/UKBB-PGS/blob/main/phenotype-info.csv

• GWAS effect sizes, ancestry-specific LD scores, and allele frequencies were computed by the Pan UKBB team, and they can be accessed at: https://pan.ukbb.broadinstitute.org/downloads

• The LD reference panel (for pruning GWAS results) and missing allele frequencies were obtained from phase 3 of 1000 Genomes Project [42]: https://www.internationalgenome.org/data-portal/data-collection/phase-3

• All code used for this paper is available at https://github.com/LachanceLab/gini/.

• Summary statistics for all 163 traits analyzed, is included as an additional file (Table S1)

## Declaration of interests

### Ethics approval and consent to participate

This research has been conducted using the UK Biobank Resource under application number 17984.

### Consent for publication

Not applicable

## Supplementary Information

Supplemental information, including three supplemental figures and one supplemental table.

