## Supplemental figures for "Different genetic architectures of complex traits and their relevance to polygenic score performance in diverse populations"

There are three supplemental figures and one supplemental table associated with this work.

**Table S1**: Summary statistics for all 163 traits analyzed in this paper (included as a separate file).

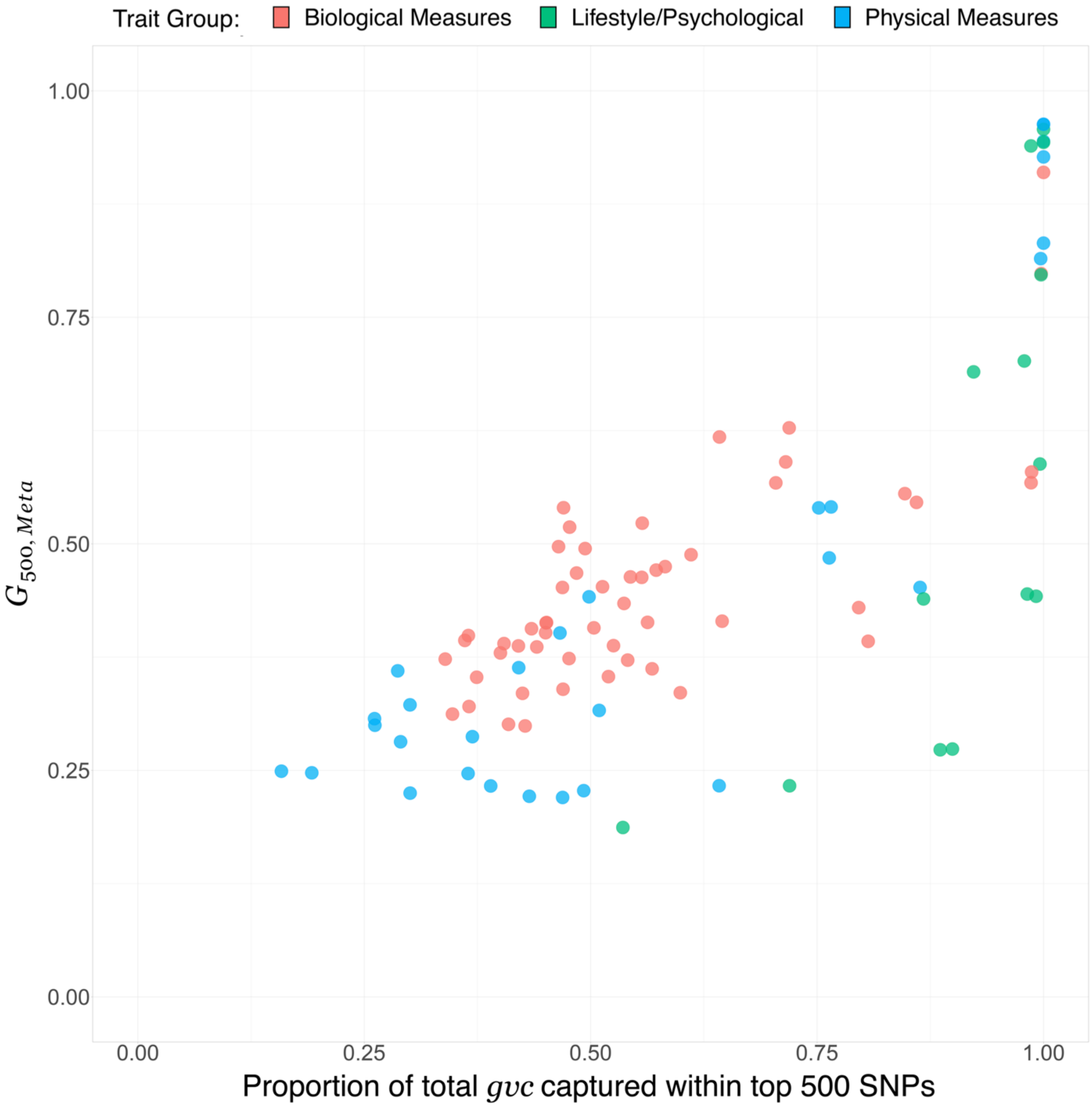


**Fig. S1.** Genomic inequality statistics (Gini) vs. the proportion of total $gvc$ captured by the top 500 SNPs. Each colored point corresponds to a different quantitative trait. The proportion of $gvc$ captured by top SNPs is the ratio of the summed $gvc$ of the top 500 SNPs to the summed $gvc$ of all independent (i.e., LD-pruned, $R^2 < 0.2$) SNPs associated with each trait (Pan UKBB meta-analysis GWAS p-value < 0.00001).

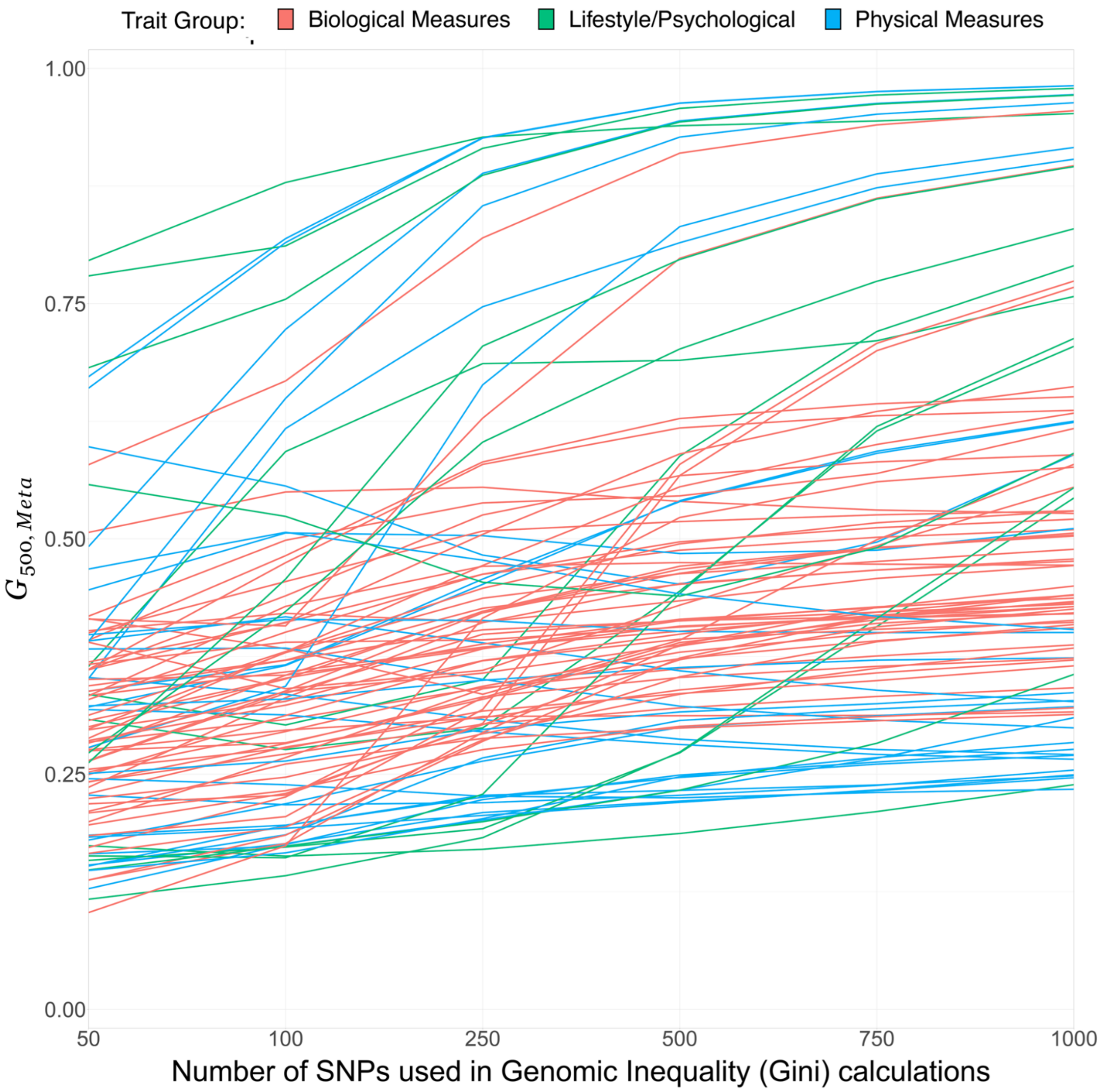


**Fig. S2.** Genomic inequality statistics (Gini) vs. the number of top SNPs used in Gini calculations. Each colored line corresponds to a different quantitative trait. As the number of SNPs increases, the Gini coefficient tends to increase as well. However, the rank orders of traits are largely conserved.

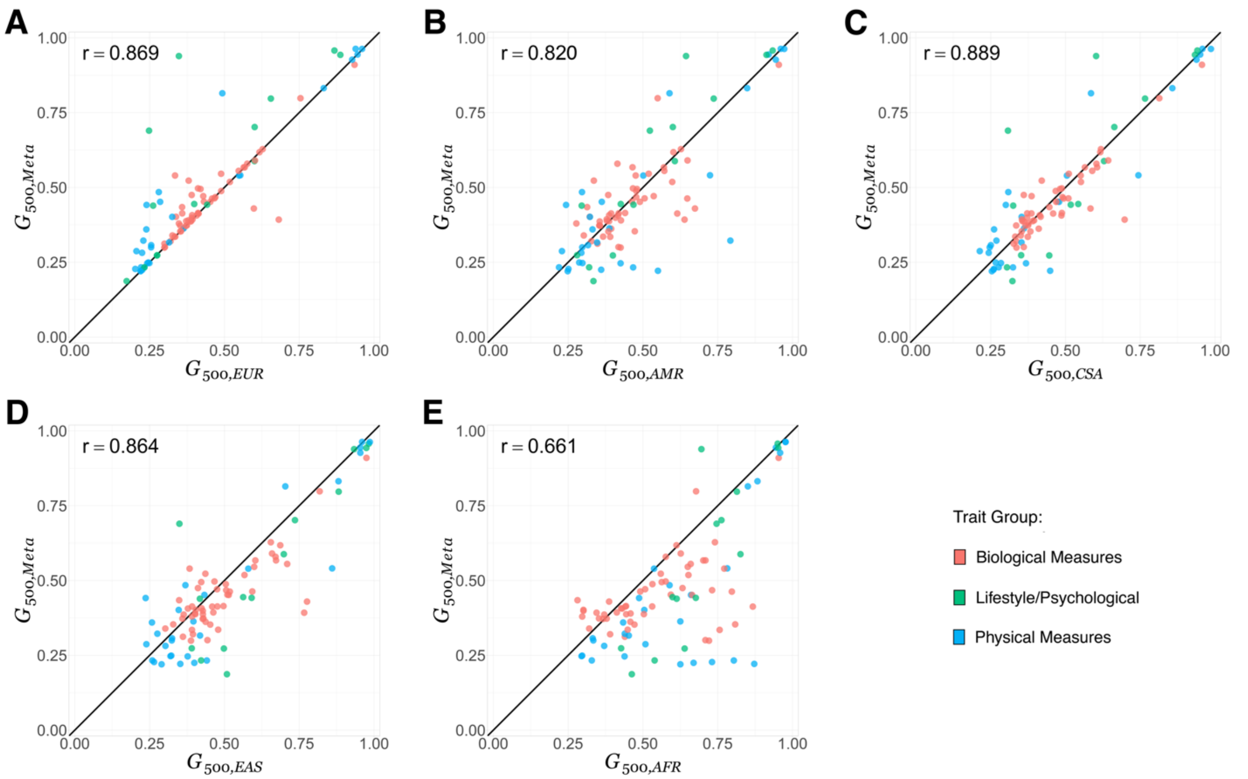


**Fig. S3.** Each panel compares the Gini coefficient computed from the meta-analysis data versus data from each of five continental ancestries: **A** European ancestry (EUR), **B** Admixed American ancestry (AMR), **C** Central/South Asian ancestry (CSA), **D** East Asian ancestry (EAS), and **E** African ancestry (AFR). All Gini coefficients were computed for the top 500 SNPs by $gvc$ within each ancestry, meaning that potentially different sets of SNPs are being considered for each ancestry within each trait. Each colored point corresponds to a different quantitative trait.
